# Uncovering genomic trajectories with heterogeneous genetic and environmental backgrounds across single-cells and populations

**DOI:** 10.1101/159913

**Authors:** Kieran Campbell, Christopher Yau

## Abstract

Pseudotime algorithms can be employed to extract latent temporal information from crosssectional data sets allowing dynamic biological processes to be studied in situations where the collection of genuine time series data is challenging or prohibitive. Computational techniques have arisen from areas such as single-cell ‘omics and in cancer modelling where pseudotime can be used to learn about cellular differentiation or tumour progression. However, methods to date typically assume homogenous genetic and environmental backgrounds, which becomes particularly limiting as datasets grow in size and complexity. As a solution to this we describe a novel statistical framework that learns pseudotime trajectories in the presence of non-homogeneous genetic, phenotypic, or environmental backgrounds. We demonstrate that this enables us to identify interactions between such factors and the underlying genomic trajectory. By applying this model to both single-cell gene expression data and population level cancer studies we show that it uncovers known and novel interaction effects between genetic and enironmental factors and the expression of genes in pathways. We provide an R implementation of our method *PhenoPath* at https://github.com/kieranrcampbell/phenopath

## Introduction

Dynamic or progressive biological behaviours are ideally studied within a longitudinal framework that allows for monitoring of individuals over time leading to direct time course data. However, longitudinal studies are often challenging to conduct and cohort sizes limited by logistical and resource availability. In contrast, cross-sectional surveys of a population are often relatively easier to conduct in large numbers and are more prevalent for molecular ‘omics based studies. Cross-sectional studies do not directly capture the changes in disease characteristics in patients but it maybe possible to recapitulate aspects of temporal variation by applying “pseudotime” computational analysis.

The objective of pseudotime analysis is to take a collection of high-dimensional molecular data from a cross-sectional cohort of individuals and to map these on to a series of one-dimensional quantities, that are called *pseudotimes.* These pseudotimes measure the relative progression of each of the individuals along the biological process of interest, e.g. disease progression, cellular development, etc., allowing us to understand the (pseudo)temporal behaviour of measured features without explicit time series data (Figure 1A). This analysis is possible when individuals in the crosssectional cohort behave asynchronously and each is at a different stage of progression. Therefore, by creating a relative ordering of the individuals, we can define a series of molecular states that constitute a *trajectory* for the process of interest.

**Figure 1:**
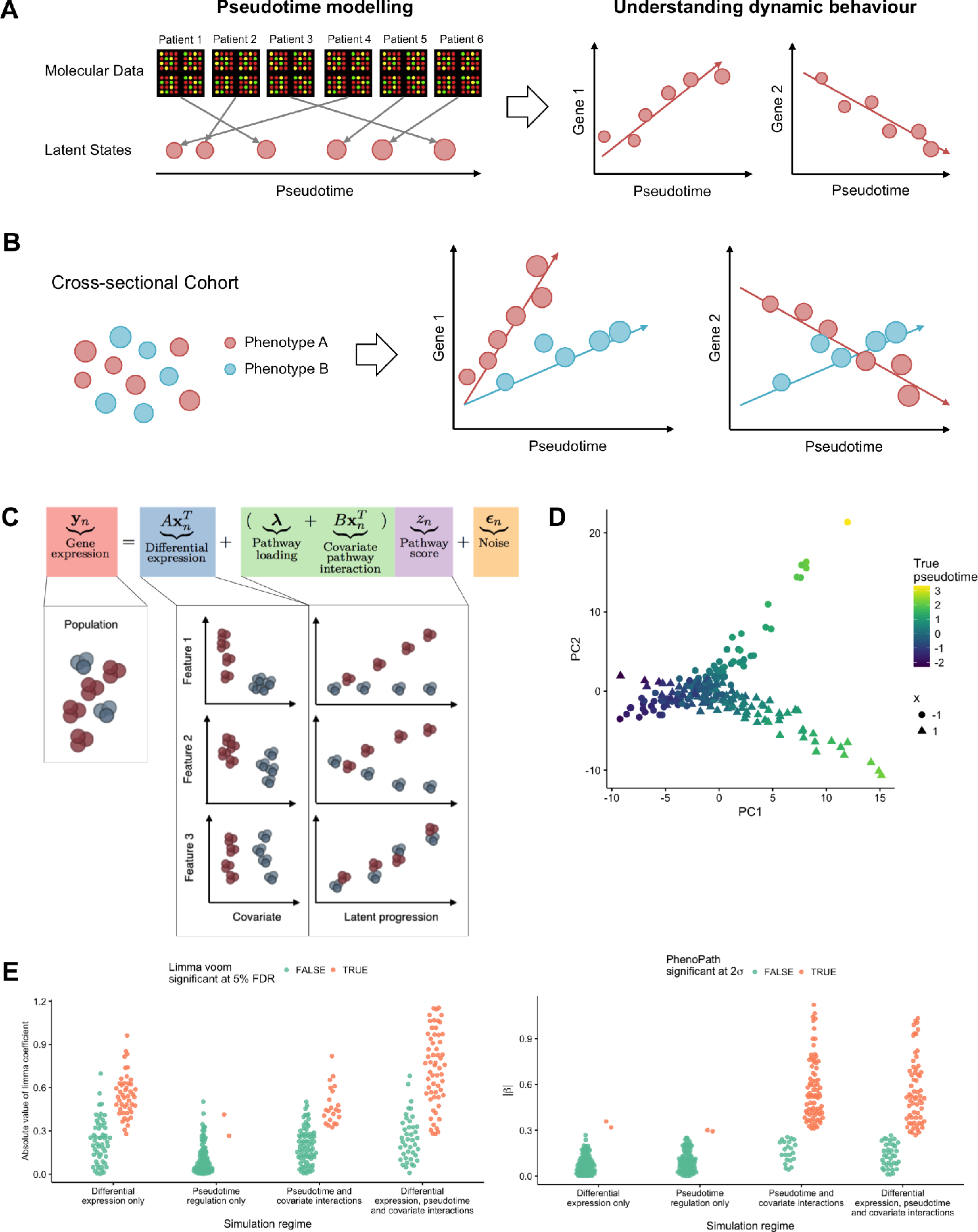
Pseudotemporal analysis. A High-dimensional molecular data from a cross-sectional cohort is mapped on to a one-dimensional pseudotemporal progression scale allowing pseudotemporal behaviour of individual features to be analysed. B If the cohort contains sub-populations we may want each sub-population to be associated to distinct trajectories. C PhenoPath models observed expression as a combination of standard differential expression and pseudotime/pathway effects, including covariate-pathway interactions. D PCA representation of a simulated dataset coloured by pseudotime shows a clear splitting of trajectories between covariate status *x* = (−1,1). E Simulation results showing the absolute value of the effect size reported by limma voom and the interaction coefficients *β* reported by PhenoPath under the four different simulation regimes and coloured by significance at 5% FDR.

Pseudotime methods generally rely on the assumption that any two individuals with similar observations should carry correspondingly similar pseudotimes and algorithms will attempt to find some ordering of the individuals that satisfies some overall global measure that best adheres to this assumption (Figure 1A). Exact implementations and specifications differ between pseudotime approaches particularly in the way “similarity” is defined and modelled. When applied to molecular data, pseudotime analysis typically captures some dominant mode of variation that corresponds to the continuous (de)activation of a set of biological pathways [Fan et al., 2016].

Pseudotime analysis has gained great popularity in the domain of single cell gene expression analysis (where each “individual” is now a single cell) in which it has been applied to model the differentiation of single-cells [Trapnell et al., 2014, Reid and Wernisch, 2016, Haghverdi et al., 2016, Campbell and Yau, 2016, Setty et al., 2016]. Using advanced machine learning techniques, these methods can be applied to characterise complex, nonlinear behaviours, such as cell cycle, and modelling branching behaviours to allow, for example, the possibility of cell fate decision making. Historically, single cell applications were pre-dated by more general applications in modelling cancer progression from gene expression profiling of tumours [Qiu et al., 2011, Magwene et al., 2003, Gupta and Bar-Joseph, 2008] as well as in other progressive disease contexts such as glaucoma [Tucker and Garway-Heath, 2010, Tucker et al., 2017, Tucker and Li, 2015, Tucker et al., 2015]. However, to date, there has been little cross-over between these domains in terms of methodological development due to the differing contexts in which methods are applied.

For instance, a limitation of pre-existing pseudotime approaches is that they generally do not provide a mechanism to account for known genetic, phenotypic and environmental information that might allow us to answer questions related to the interaction between heterogeneity in these factors and pseudotime progression. For example, do immune cells exposed to different stimuli progress differently in their response? Do the transcriptional programmes of tumours differ based on mutation status of a known cancer gene? Whilst pseudotime methods exist for unsupervised identification of multiple or branching pseudotime trajectories, these can only be *retrospectively* examined for their association with external factors of interest and do not provide an explicit approach for identifying associations.

In this paper, we describe a novel Bayesian statistical framework for pseudotime trajectory modelling to address these limitations. Our framework models global pseudotemporal progression but incorporates covariates that can modulate the pseudotemporal progression allowing sub-groups within the cross-sectional population to each develop their own trajectory (Figure 1B). Our approach combines linear regression and latent variable modelling approaches and allows for interactions between the covariate and temporally driven components of the model. We believe our method to be the first integrated statistical approach for modelling pseudotime trajectories against heterogeneous genetic and environmental backgrounds allowing its utility in both single and non-single cell applications.

## Results

### PhenoPath: a Bayesian statistical framework for learning continuous pathway or pseudotemporal with covariates

We first give an overview of our statistical method which we call “PhenoPath”. For simplicity, our descriptions will assume that the observed data are high-dimensional gene expression measurements which are used throughout our empirical experiments but we stress that the model would be applicable to a wider range of data modalities. PhenoPath uses a Bayesian statistical framework that combines linear regression and latent variable modelling. The observed data (**y**_n_) for the *n*-th individual is a linear function of both measured covariates (**x**_n_) and an unobserved latent variable (*z_n_*) corresponding to pseudotime. We will also refer to this latter quantity more generically as a *pathway score* since, as we will explore further, pseudotime progression will be driven by the activities of certain biological pathways. Figure 1C shows a schematic of the model. The covariate-dependent component (A**x**_n_) models differential expression whilst the pseudotime component involves both a pathway-only component (***λ***). The key novelty is an interaction term 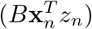 that allows the the covariates to modulate the pathway or pseudotemporal trajectory. We devise a Bayesian hypothesis test for the model to test for these interaction effects (see Methods for details). An attractive feature of the framework is that, in the absence of pseudotemporal variation, the model reduces to a standard differential expression model. Whilst, in the absence of measured covariates, it is a factor analysis latent variable model for pseudotime.

In our investigations, the covariates will be discrete, binary quantities but this is not a necessary restriction. Sparse Bayesian prior probability distributions are used to constrain the parameters (A, B, λ) so that covariates only drive the emergence of distinct trajectories only if there is sufficient information within the data to do so. Computational inference within PhenoPath is handled by a fast and highly scalable variational Bayesian inference framework that can handle thousands of features and samples in minutes using a standard personal computer making it readily applicable to large data sets without the use of high-performance computing (see Methods for details).

### Simulation study

We first demonstrate the utility of our model by performing a simulation study to demonstrate the value of modelling covariate-pathway interactions. We simulated RNAseq-based gene expression data [Frazee et al., 2015] where genes were either (1) differentially expressed only, (2) exhibiting pseudotime progression only, (3) driven by covariate-modulated pseudotime progression, or (4) differentially expressed with covariate-modulated pseudotime progression (see Methods for details, Figure 1D, Supplementary Fig. 1). PhenoPath exhibited high specificity and sensitivity by classifying only a small number of simulated genes (2%) as exhibiting interaction effects in cases 1-2 where there are no covariate-pseudotime interactions but identifies 78% and 63% of genes as exhibiting significant covariate-pseudotime interactions in cases 3 and 4 respectively (Fig 1E). For comparison, a standard differential expression analysis using limma-voom identified 47% and 59% of genes as differentially expressed in cases 1 and 4 respectively. In case 2 only 2% of genes are identified as DE as expected but, in case 3, 22% of genes are identified as DE where limma-voom would not be expected to report any differentially expressed genes. e sought to compare the performance of Limma Voom and PhenoPath in detecting differential expression and pathway interaction effects respectively, and show that there are pathway interaction effects not evident from differential expression analyses alone. We found that PhenoPath identifies such interactions with high precision (Supplementary Table 1).

### Single-cell RNA-seq perturbation analysis

We next examined a time-series single-cell RNA-seq (scRNA-seq) data set of bone marrow derived dendritic cells responding to particular stimuli [Shalek et al., 2014]. Cells were exposed to LPS, a component of Gram-negative bacteria, and PAM, a synthetic mimic of bacterial lipopeptides, and scRNA-seq performed at 0, 1, 2, 4 and 6 hours after stimulation. Despite the time-series measurement, previous studies have suggested this dataset is more suited to a “pseudotime” analysis as the cells respond asynchronously and heterogeneity exists within the cellular populations at each time point [Reid and Wernisch, 2016]. To-date pseudotime inference algorithms would typically assume a common trajectory across all experimental conditions or a pseudotime analysis performed separately for each stimulant. This might give a loss of statistical power and artefacts introduced by confounding effects. Using our model we can encode the stimulant to which the cells were exposed as a covariate and allow gene expression to evolve along pseudotime differently for either LPS or PAM exposure. This allows us to learn a single trajectory for all cells regardless of stimulant applied yet simultaneously infer which genes are differentially regulated in response. We applied this to the 820 cells exposed to LPS and PAM in the time points 1, 2, 4, and 6 hours after stimulation using the 7,533 genes whose variance in normalised log-expression exceeded a pre-set threshold (see Methods for details).

We inferred a covariate-perturbed trajectory and uncovered a landscape of pseudotime-stimulant interactions (Fig 2A), unveiling genes whose regulation along pseudotime is modulated by the application of LPS or PAM. The trajectory inferred largely recapitulated the true time-series measurement (Fig 2B, R^2^ = 0.64), despite no explicit temporal information being provided to the algorithm, though transcriptional heterogeneity at each time point is still evident. We also compared this to two commonly-used pseudotime algorithms and found that the pseudotimes inferred using PhenoPath had the best agreement with the capture times (Supplementary Fig. 2), possibly due to the ability to integrate the confounding effect of differential stimulant exposure.

**Figure 2.**
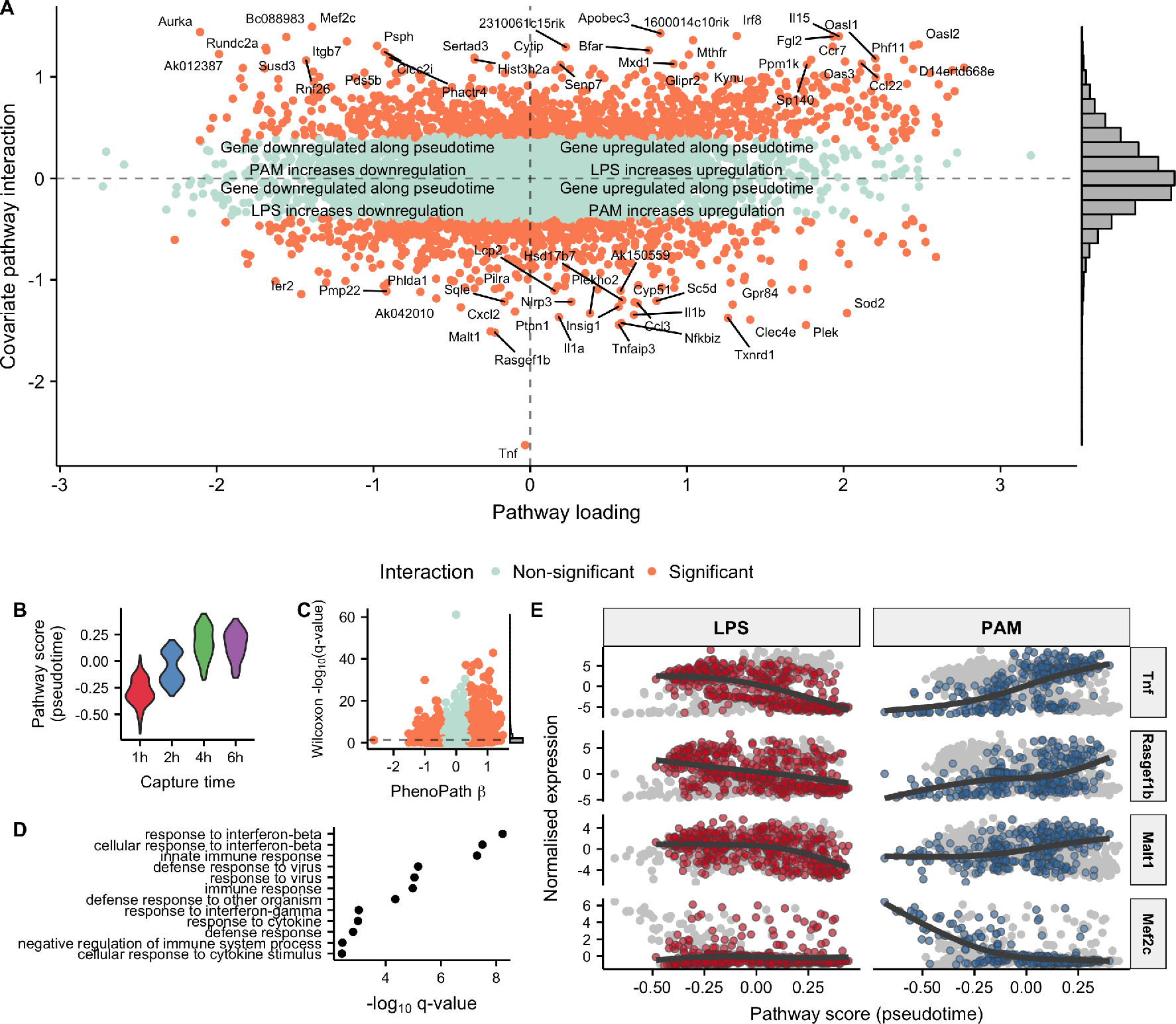
Stimulant-immune reactions in single-cell RNA-sequencing data. **A** PhenoPath applied to the Shalek et al. dataset uncovers genes differentially regulated along pseudotime depending on the stimulant (LPS or PAM) applied. **B** The inferred pseudotimes (*z*) consistent with the physical capture times. **C** A comparison of *p*-values obtained through a nonparametric statistical test for differential expression between LPS and PAM stimulation shows no particular relation with the interaction parameters *β* inferred with PhenoPath. **D** A GO enrichment analysis of the genes upregulated along pseudotime whose upregulation was increased by LPS stimulation showed enrichment for immune system processes. **E** Expression of the four genes with the largest interaction effect sizes along over pseudotime, stratified by stimulant applied. Strikingly, *Tnf* is upregulated under PAM exposure yet downregulated under LPS stimulation.

Using PhenoPath we discovered a large number of stimulant-modulated interactions masked by standard differential-expression analysis (Fig 2C). A GO analysis revealed genes whose upregulation along the common trajectory was increased by LPS exposure (as opposed to PAM) were highly enriched for interferon-beta and immune response (Fig 2D), which recapitulates previous results [Shalek et al., 2014, Reid and Wernisch, 2016] that suggest a “core” module of antiviral genes upregulated at later timepoints in LPS cells but in an entirely unsupervised, integrated manner. We finally examined the individual genes most perturbed by LPS or PAM along the trajectory (Fig 2E), which identifies as yet uncharacterised expression patterns associated with LPS and PAM. Most notably, the tumour necrosis factor Tnf had around twice the interaction effect size of any other gene, and decreases under LPS stimulation but increases under PAM. Further genes exhibit differential regulation according to stimulant, such as *Mef2c* that has constant expression over pseudotime under LPS stimulation yet shows downregulation under PAM stimulation. These results complement previously discovered gene differences such as that of *Tnf*, but in a systematic, transcriptome-wide approach.

### Identifying microsatellite instability associated gene expression heterogeneity in colorectal cancer

We next applied our model to a non-single cell setting by examining RNA-seq gene expression data from the TCGA colorectal adenocarcinoma (COAD) cohort [Network et al., 2012]. We used microsatellite instability status (MSI) as a phenotypic covariate and wanted to identify pseudotemporal expression patterns associated with MSI status. MSI is genetic hypermutability that is present in around 15% of colorectal tumours and is associated with differential response to chemotherapeutics and marginally improved prognosis [Boland and Goel, 2010].

We applied PhenoPath to 4,801 highly variable genes across 284 samples to identify a pseudotemporal trajectory through these tumours (see Methods for details). This analysis uncovered a landscape of 92 pathway-MSI interactions including known tumour suppressor genes (Fig 3A & Supplementary Data 1). Patients further advanced along the trajectory exhibited higher expression of T regulatory cell (Tregs) immune markers (Fig 3B) likely due to increasing T regulatory cell infiltration of the tumour. This led us to hypothesise that the inferred pseudotime trajectory corresponds to immune response pathway activation in the tumours, further supported by a Gene Ontology (GO) enrichment analysis for genes upregulated along the trajectory (Fig 3C). Tumour-infiltrating Tregs are potent immunosuppressive cells of the immune system that promote progression of cancer through their ability to limit antitumuor immunity and promote angiogenesis and often associated with a poor clinical outcome [Facciabene et al., 2012]. A standard differential expression analysis using limma voom [Law et al., 2014] (Fig 3D) demonstrates that PhenoPath is required to uncover such interactions as a gene being differentially expressed does not imply a pathway-MSI interaction, while such interactions do not require differential expression.

**Figure 3:**
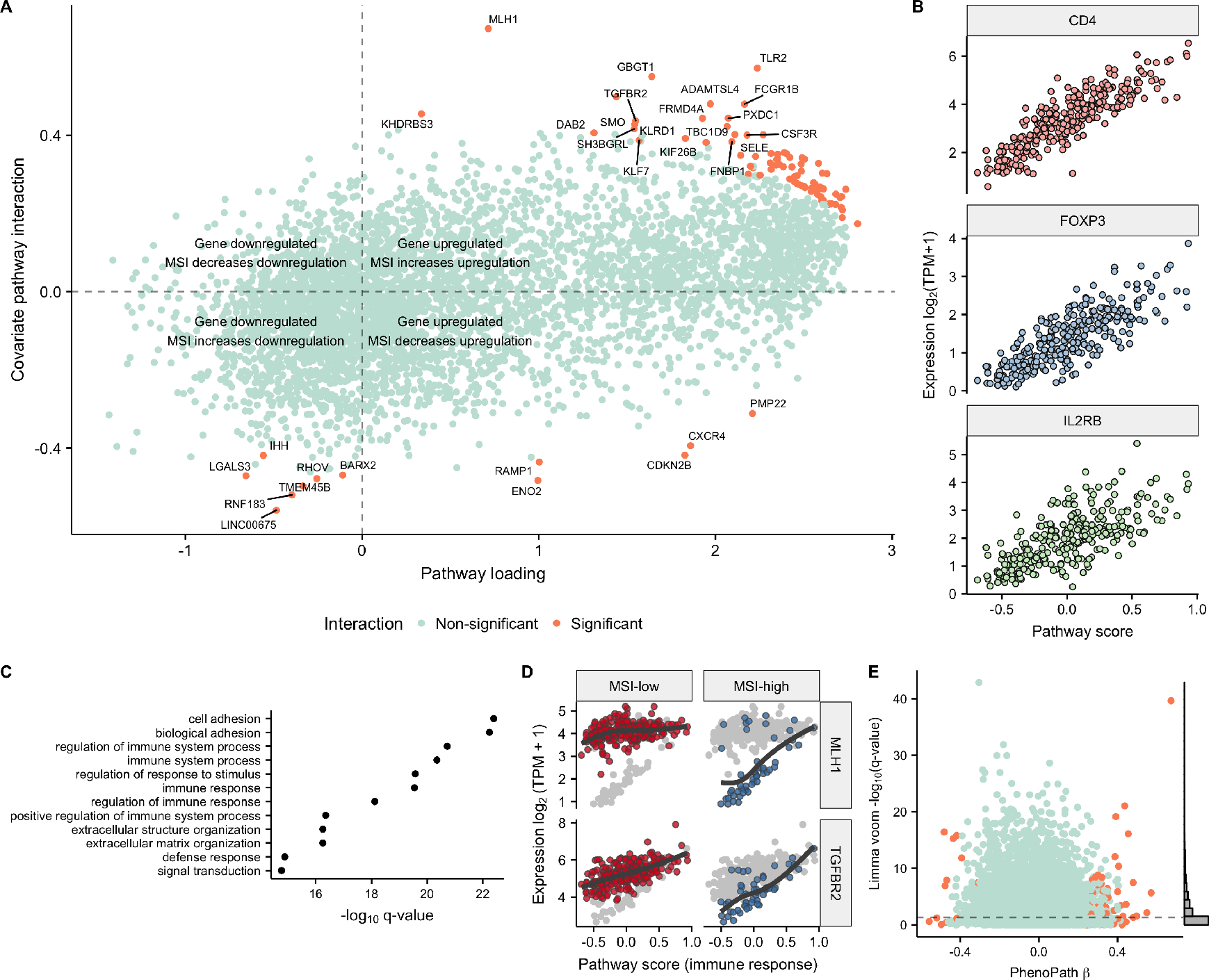
Immune-microsatellite instability interactions uncovered in colorectal adenocarcinoma. **A** PhenoPath applied to colorectal adenocarcinoma (COAD) RNA-seq expression data uncovers a landscape of interactions between the inferred immune trajectory and microsatellite instability status (MSI). **B** Expression of three T regulatory cell markers demonstrates that our pseudotime corresponds to activation of immune response pathways. **C** A comparison to the FDR-corrected *q*-values reported by Limma Voom demonstrates genes found interacting with MSI status and the immune pathway are found to be both DE and non-DE in standard analyses. **D** A GO enrichment analysis of upregulated genes implies the latent trajectory encodes immune pathway activation in each tumour. **E** The tumour suppressor genes *MLH1* and *TGFBR2* were identified by our method as being significantly perturbed along the immune trajectory by MSI status. *MLH1* shows no interaction with immune pathway activation in the MSI-low regime yet is highly correlated with immune pathway activation in the MSI-high regime.

The most striking interaction discovered for this dataset was the *MLH1* gene whose interaction effect size was far larger than any other gene. This association provided a positive control since *MLH1* is a DNA mismatch repair gene and germline mutations of which are causal for hereditary non-polyposis colorectal cancer [Bonadona et al., 2011, Gille et al., 2002]. By applying PhenoPath we correctly identified that in patients, with low or absent levels of microsatellite instability, there is no relationship between *MLH1* expression and immune pathway interaction, with *MLH1* expressed at an approximately constant level (Fig 3E). However, when MSI occurs in a tumour, *MLH1* expression is highly correlated with immune response, showing almost no expression when the immune pathway is inactive and gradually being upregulated with immune pathway response [Michel et al., 2008].

### Tracking ER modulated angiogenesis driven progression in breast cancer

We next performed a pseudotemporal analysis of the TCGA breast cancer cohort using estrogen receptor (ER) status as a phenotypic covariate. Approximately 60% of breast cancers are estrogen receptor positive [Early Breast Cancer Trialists’ Collaborative Group (EBCTCG)], which is typically associated with improved prognosis and a longer time to recurrence [Parl et al., 1984]. We applied PhenoPath to 1,135 samples post-QC and 4,579 highly variable genes (see Methods for details). Using stringent significance testing threshold we found 1,932 genes (42%) affected by an interaction between the pseudotemporal trajectory and ER receptor status (Fig 4E & Supplementary Data 2). There was a correlation between the pathway interaction strength and the p-value reported through standard differential expression (Fig 4B), though there remained some genes that exhibited pathway interaction and no differential expression.

**Figure 4:**
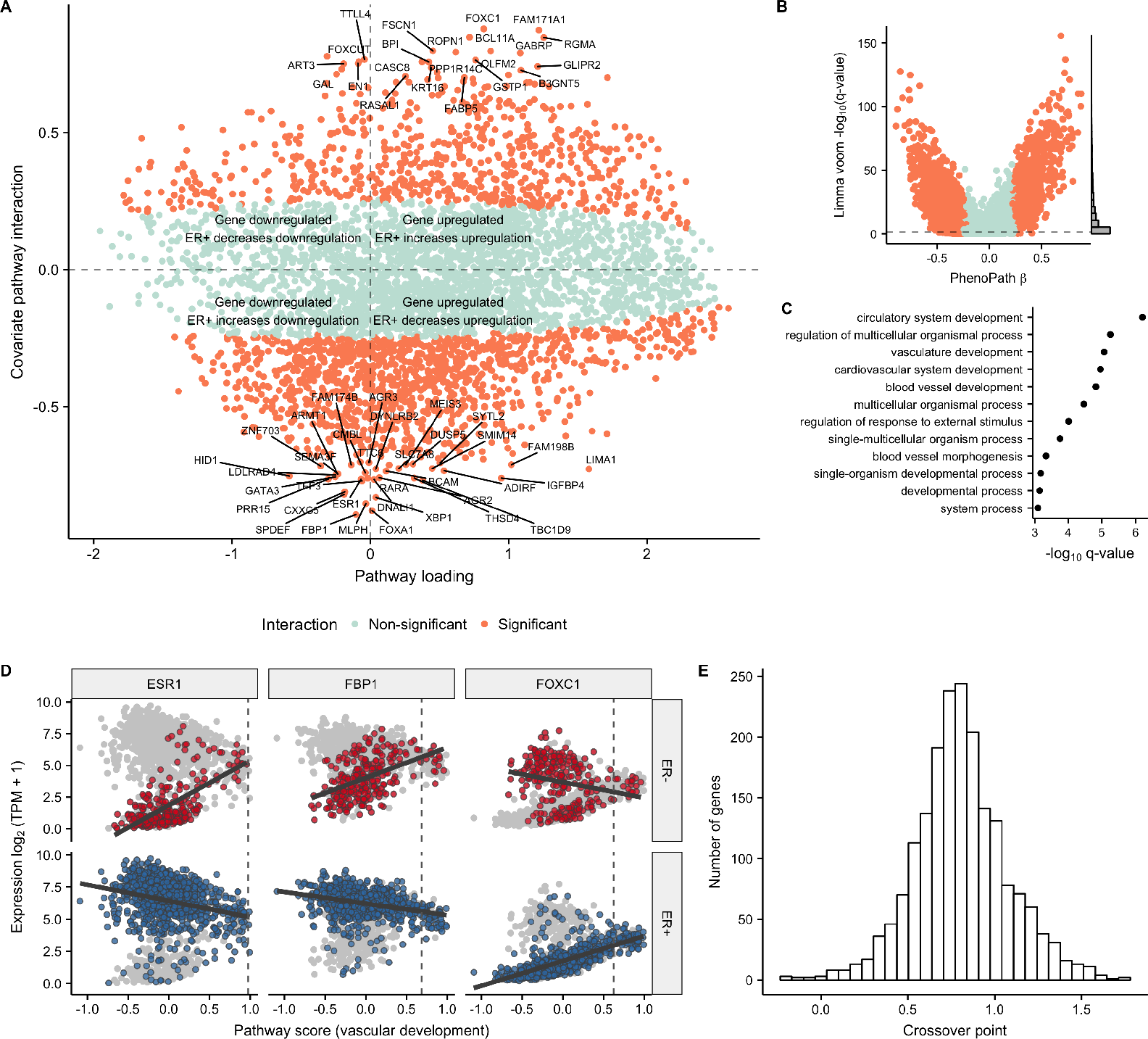
Vascular growth-ER status interactions uncovered by PhenoPath in breast cancer. **A** PhenoPath applied to Breast Cancer (BRCA) RNA-seq expression data uncovers a landscape of interactions between the inferred angiogenesis trajectory and estrogen receptor (ER) status. **B** A comparison to the FDR-corrected q-values reported by Limma Voom identifies a significant number of DE genes display an interaction with ER status and the angiogenic pathway. **C** A GO enrichment analysis of upregulated genes implies the latent trajectory encodes angiogenesis pathway activation in each tumour. **D** Four example genes *ESR1*, *FBP1*, and *FOXC1* were identified by PhenoPath as significantly perturbed along the angiogenesis trajectory by ER status. The vertical dashed line signifies the calculated crossover point, demonstrating the expression profiles of these genes converge towards the end of the trajectory. **E** A histogram of the crossover points of all genes whose trajectory-covariate interactions were significant. The vast majority of crossover points are at the end of the trajectory (around 0.5, where the “middle” pathway score is 0) implying a convergence of gene expression as the trajectory progresses.

A GO enrichment analysis indicated that the inferred pseudotemporal trajectory corresponded to vascular growth pathways or *angiogenesis* (Fig 4F) - a well-known and uncontroversial hallmark of cancer development [Ferrara, 2002, Welti et al., 2013]. We confirmed this finding by specifically examining the expression of known angiogenesis inducing genes (Supplementary Fig. 4). We found increasing fibroblast growth factor-2 (*FGF-2*) and vascular endothelial growth factors C and D (*VEGF-C/D*) expression along the trajectory whose behaviours were independent of ER status.

We finally sought to examine the genes identified as being most affected by the interaction between angiogenesis and estrogen receptor status. Importantly, this set included the Estrogen Receptor 1 (*ESR1*) gene as well as the forkhead transcription factors *FOXA1* and *FOXC1* which are known to be involved with ERα mediated action in breast cancer [Lam et al., 2013, Yu-Rice et al., 2016] (Fig 4D and Supplementary Fig. 3). Fig 4D shows how the fructose-1,6-biphosphatase (*FBP1*) and *FOXC1* genes evolve along the angiogenesis pathway dependent on ER status. In the ER-regime, *FBP1* is upregulated along the trajectory while in the ER+ regime it is downregulated. Intriguingly, *FBP1* has been identified as a marker to distinguish ER+ from ER-subtypes and its expression has been shown to be negatively correlated with *SNAIL* as the Snail-G9a-Dnmt1 complex, is critical for E-cadherin promoter silencing, and required for the promoter methylation of FBP1 in basal-like breast cancer [Dong et al., 2013] (Supplementary Fig. 5). Similarly, *FOXC1* shows no regulation in the ER-regime yet is strongly upregulated in the ER+ case.

We noted that these genes exhibit a convergence - they have markedly different expression at the beginning of the trajectory based on ER status yet converge towards the end. We derived a mathematical formula to infer such convergence points and calculated these for all genes showing significant interactions (see Methods for details). Remarkably, the vast majority converge towards the end of the trajectory (Fig 4E), implying a common end-point in vascular development for both ER+ and ER-cancer subtypes (Supplementary Fig. 6). This effect can be seen in the example expression plots in Figure 4D, where the vertical dashed line represents the convergence point always at the end of the trajectory. This suggests that while there exists low levels of angiogenesis pathway activation, ER status dominates gene expression while as angiogenesis pathway activation increases it comes to dominate expression patterns over ER status. This finding might have implications for the application of angiogenesis inhibitors in breast cancer treatment.

## Discussion

PhenoPath provides a novel contribution to the pre-existing arsenal of pseudotemporal analysis algorithms developed across a range of application areas including single cell ’omics and cancer. Using a statistical model that allows for covariate-modulated pseudotemporal trajectories, PhenoPath generalises pseudotime analysis to a wider range of applications where genetic, phenotypic or environmental contexts may vary between samples and be influential in the trajectories. We have demonstrated its utility in an application to single cell transcriptomics involving external stimuli and there is potential usage in high-throughput single cell CRISPR experiments that are as yet unexplored [Adamson et al., 2016, Datlinger et al., 2017]. We also demonstrated applications to The Cancer Genome Atlas using PhenoPath to model disease trajectories in colorectal and breast cancer. The trajectories identified were consistent are consistent with pre-existing knowledge concerning tumorigenesis in these disease. Importantly, PhenoPath was able to identify covariate-pathway interactions that might be driving specific trajectory differences recovering known associations as well as novel genes. We showed that these behaviours cannot be readily determine with standard differential expression analyses without taking into account the latent disease progression. In summary, PhenoPath provides a powerful and scalable pseudotime analysis algorithm for modelling latent progression in a variety of experimental settings. Future work will expand the ability of PhenoPath to handle complex mixtures of continuous and discrete covariates in high-dimensional settings.

## Methods

### Statistical model

We begin with an *N* × *G* data matrix **Y** where *y_ng_* denotes the *n^th^* entry in the *g^th^* column for *n* ∈ 1,…, N samples and *g* ∈ 1,…, *G* features. Such a matrix would correspond to the measurement of a dynamic molecular process that we might reasonably expect to show continuous evolution such as gene expression corresponding to a particular pathway. It is then trivial to learn a one-dimensional linear embedding that would be our “best guess” of such progression via a factor analysis model:

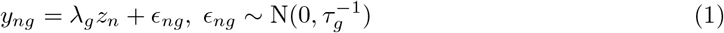

where *z_n_* is the latent measure of progression for sample *n* and λ_g_ is the factor loading for feature *g* which essentially describes the evolution of *g* along the trajectory.

However, it is conceivable that the evolution of feature *g* along the trajectory is not identical for all samples but is instead affected by a set of external covariates. Note that we expect such features to be “static” and should not correlate with the trajectory itself.

Introducing the *N × P* covariate matrix **X** with the entry in the *n^th^* row and *p^th^* column given by *x_np_,* we allow such measurements to perturb the factor loading matrix

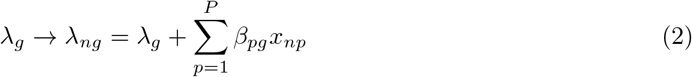

where *β_pg_* quantifies the effect of covariate *p* on the evolution of feature *g*. Despite **Y** being column-centred we need to reintroduce gene and covariate specific intercepts to satisfy the model assumptions, giving a generative model of the form

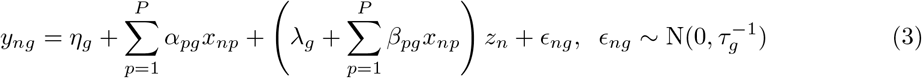

Our goal is inference of *z_n_* that encodes progression along with *β_pg_* which is informative of novel interactions between continuous trajectories and external covariates. Consequently we place a sparse Bayesian prior on *β_pg_* of the form 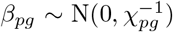 where the posterior of *χ_pg_* is informative of the model’s belief that *β_pg_* is non-zero. The complete generative model is therefore given by

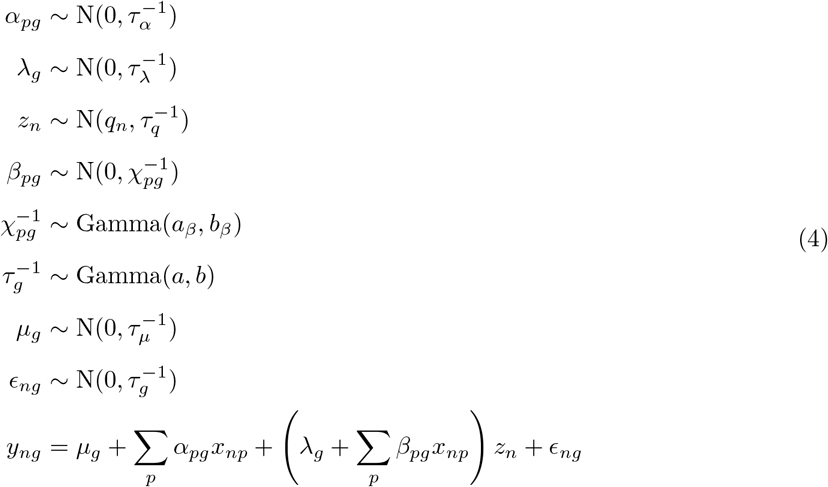

where *T*_α_, *T*_λ_, *a*, *b*, *a_β_*, *b_β_*, *τ_q_* are fixed hyperparameters and *q_n_* encodes prior information about *z_n_* if available but typically *q_n_* = 0 ∀*i* in the uninformative case.

To understand this model it helps to consider the distribution of **Y** marginalised over the mapping {*λ_g_,α_*pg*_, β_*pg*_*} ∀ *p*, *g* with priors 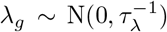 and 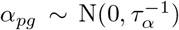. If **y**_g_ denotes the column vectors of **Y** and similarly **x***_p_* for **X**, [**z**]*_n_* = *z_n_*, **1**_N_ is the column vector of ones and ⊙ denotes the element-wise product, then

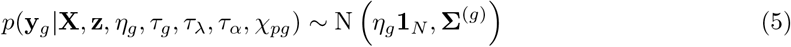

where

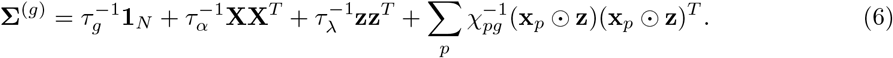

We therefore see that the addition of the covariates adds extra terms to the covariance matrix corresponding to *perturbations* of the latent variables with the covariates. Consequently, the scale on which ***x****_p_* is defined needs carefully calibrated. Furthermore, it is possible to extend the latent variable matrix to have dimension larger than 1 giving a novel dimensionality reduction technique for visualisation, though additional rotation issues arise.

### Inference

We perform co-ordinate ascent mean field variational inference (see e.g. [Blei et al., 2016]) with an approximating distribution of the form

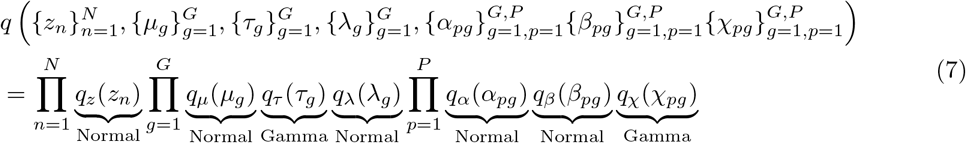

Due to the model’s conjugacy the optimal update for each parameter *θ_j_* given all other parameters *θ_−j_* can easily be computed via

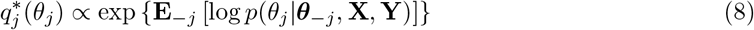

where the expectation is taken with respect to the variational density over θ_−j_.

### Identifying significant interactions

For each gene *g* and covariate *p* we have *β_pg_* that encodes the effect of *p* on the evolution of *g* along the trajectory **z**. We would like to identify interesting or *significant* interactions for further analysis and follow up.

The variational approximation for *β_pg_* is given by

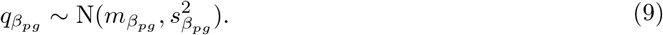

We therefore define an interaction as significant if 0 falls outside the posterior *nσ* interval of *mβ_pg_*. In other words, the interaction between *p* and *g* is significant if

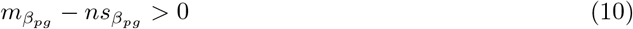

*or*

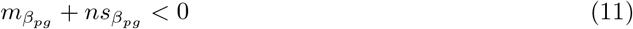

Note that variational inference typically underestimates posterior variances [Blei et al., 2016] so such a designation of *significant* will be under-conservative. For all analyses we select *n* = 3, which would loosely correspond to 0 being outside the 99.7% posterior interval of *β_pg_*.

### Synthetic data study

We performed a small simulation study to identify effects uncovered by PhenoPath that are missed by standard differential expression analyses. Specifically, we sought to compare differentially expressed genes identified by limma voom [Law et al., 2014], one of the leading RNA-seq differential expression methods, to the *β* interactions from PhenoPath. For *N* = 200 samples we assigned each to one of two categories given by the *x* values *x* = −1,1, and assigned a pseudotime *z* through draws from a standard normal distribution. For each sample *i* = 1,…, *N* and gene *g* = *1,…,G* we then generated a mean value through the PhenoPath mean function

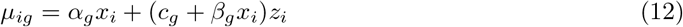

The gene-specific parameters (*α_g_*, *c_g_*, *β_g_*) were sampled in equal proportions from one of four classes:

1. *Differential expression only* where *α_g_* = 1 or −1 with equal probability and *c_g_* = *β_g_* =0
2. *Pseudotime regulation only* where *c_g_* = 1 or −1 with equal probability and *α_g_* = *β_g_* =0
3. *Pseudotime and covariate interactions* where *c_g_* and *β_g_* are set to 1 or −1 with equal probability and *α_g_* = 0
4. *Differential expression, pseudotime and covariate interactions* where all parameters take on values of −1 or 1 with equal probabilities

In order to generate RNA-seq reads we need positive count values. In the spirit of general linear models, we then used *g*(*x*) = 2^*x*^ as a link function and generated a matrix of positive means

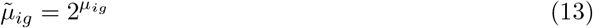

We subsequently simulated a count matrix *c_ig_* by sampling for each entry from a negative binomial distribution with mean 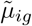 and size parameter 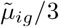. While this could be used as input to PhenoPath (suitable log transformed), we sought to make our simulation as realistic as possible including quantification errors. We subsequently simulated FASTA files using the Bioconductor package `polyester` [Frazee et al., 2015] using the first 400 transcripts of the reference transcriptome of the 22nd human chromosome. FASTA files were then converted to FASTQ files using a script copied from StackOverflow and quantified into TPM and count estimates using Kallisto [Bray et al., 2016]. The log_2_(TPM +1) values were then used for input to PhenoPath while the raw count values were used for input to limma voom.

In our simulation study, Limma Voom “only” detects 47% of the genes simulated as differentially expressed. Such power to detect differential expression is dependent on effect sizes and measurement noise, and so such a figure is in no way unreasonable given the parameters used. While a more comprehensive simulation study could examine detection rates across entire distributions over effect sizes and measurement noise, we simply sought to perform a simulation that demonstrated that PhenoPath identifies a subset of differential expression and that standard differential expression misses some interactions across a consistent effect size and noise regime.

### Fitting pseudotimes to Shalek et al. dataset

The Shalek et al. dataset of time-series dendritic cells was previously used in a pseudotime analysis where the capture times were explicitly used as priors on the latent space [Reid and Wernisch, 2016]. However, in PhenoPath we provide no explicit temporal information, so sought to perform a brief comparison to two popular pseudotime algorithms, Monocle 2 [Qiu et al., 2017] and DPT [Haghverdi et al., 2016]. For both methods we provided the same normalised log expression (see section below) and ran the algorithms with the default parameters. Performance of each algorithm was assessed by regressing the inferred pseudotimes on the capture times using the `R` function `lm` and computing the *R^2^.*

### Data retrieval and processing

#### Shalek et al

Preprocessed TPM values for all cells were retrieved from the Gene Expression Omnibus (`GSE48968`). We retained cells treated by LPS and PAM at time points 1h, 2h, 4h, and 6h, resulting in 820 cells (479 LPS and 341 PAM). We retained the 7533 genes whose variance in log_2_(TPM + 1) expression was greater than 2. The first principal component of the data showed a strong dependency on the number of features expressed - previously been implicated in technical effects [Hicks et al., 2015] - which we subsequently removed using the `normalizeExprs` function in `Scater` [McCarthy et al., 2017].

#### TCGA studies

For both COAD and BRCA studies, TPM matrices were retrieved from a recent transcript-level quantification of the entire TCGA study [Tatlow and Piccolo, 2016]. Clinical metadata, including the phenotypic covariates used in PhenoPath, were retrieved using the RTCGA `R` package [Kosinski and Biecek, 2016]. Transcript level expression estimates were combined to gene level expression estimates using `Scater` [McCarthy et al., 2017].

### Quality control and removal of samples

#### COAD

A PCA visualisation of the COAD dataset showed two distinct clusters based on the plate of sequencing. Rather than try to correct such a large batch effect, we retained samples with a PC1 score of less than 0 and a PC3 score greater than −10, and removed any “normal” tumour types. For input to PhenoPath we used the 4801 genes whose median absolute deviation in log(TPM + 1) expression was greater than 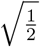.

#### BRCA

A PCA visualisation of the BRCA daatset showed a loosely dispersed outlier population that separated on the first and third principal components. We performed Gaussian mixture model clustering using the `R` package `mclust`[Fraley et al.], and removed samples designated as cluster 2 in the PCA plot, giving 1135 samples for analysis. For input to PhenoPath we used the 4579 genes whose variance in log(TPM+ 1) expression was greater than 1 and whose median absolute deviation was greater than 0.

### Identifying crossover points in BRCA

In PhenoPath we model gene expression evolving along the trajectories separately for each phenotype (or covariate) considered. Unless the gradient of change along the trajectory is exactly equal for both phenotypes (i.e. *β* = 0 exactly), the gene expression will cross at a given point in the trajectory.

Inference of this point would allow us to identify sections of the trajectory not affected by the covariate and consequently sections of the trajectory that are. This is important as if the crossover point occurs towards the beginning of the trajectory, it would mean gene expression is similar at the beginning but diverges as we move along the trajectory. Similarly, if the crossover points occur towards the end of the trajectory, it would imply the expression profiles for the two phenotypes are different at the beginning of the trajectory, but converge as the trajectory progresses. An interpretation of this would be that the effect on expression from the trajectory slowly dominates over the effect of phenotypes on the trajectory.

It is important to note that the latent trajectory values loosely follow a N(0,1) distribution. This means the ‘middle’ of the trajectory is any value around zero, values of −1 or less could be thought of as the ‘beginning’ while values greater than 1 may be thought of as the ‘end’. Crucially, we can derive an analytical expression from the PhenoPath parameters for the crossover point *z*^*^ (see below).

We fitted the crossover points *z*^*^ for all *significant* genes in the BRCA dataset. We find that the vast majority of the crossover times *z*^*^ occur towards the end of the trajectory, with a median i value of around 0.4. In other words, at the beginning of the trajectory most genes are differentially expressed based on ER status, while as the trajectory progresses it comes to dominate at the gene expression converges.

### Inference of convergence point

The condition for the crossover point is that the predicted expression for each phenotype is identical. Therefore (in the context of BRCA cancer)

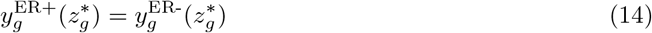

which leads to the condition

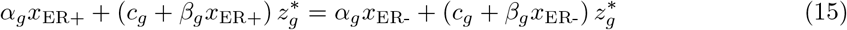

which is in turn solved by

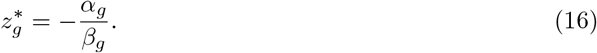

## Acknowledgements

K.C. is supported by a UK Medical Research Council funded doctoral studentship. C.Y. is supported by a UK Medical Research Council New Investigator Research Grant (Ref. No. MR/L001411/1) and Methodology Research Grant (MR/P02646X/1) and the Wellcome Trust Core Award Grant Number 090532/Z/09/Z.

---

**Supplementary Table 1:** A comparison of true positive, false positive, and false discovery rates for Limma Voom detecting differential expression and PhenoPath detecting covariate-pseudotime interactions on synthetic data.

| Algorithm | True positive rate | False positive rate | False discovery rate |
| --- | --- | --- | --- |
| Limma Voom | 0.82 | 0.09 | 0.18 |
| PhenoPath | 0.97 | 0.02 | 0.03 |

**Supplementary Figure 1:**
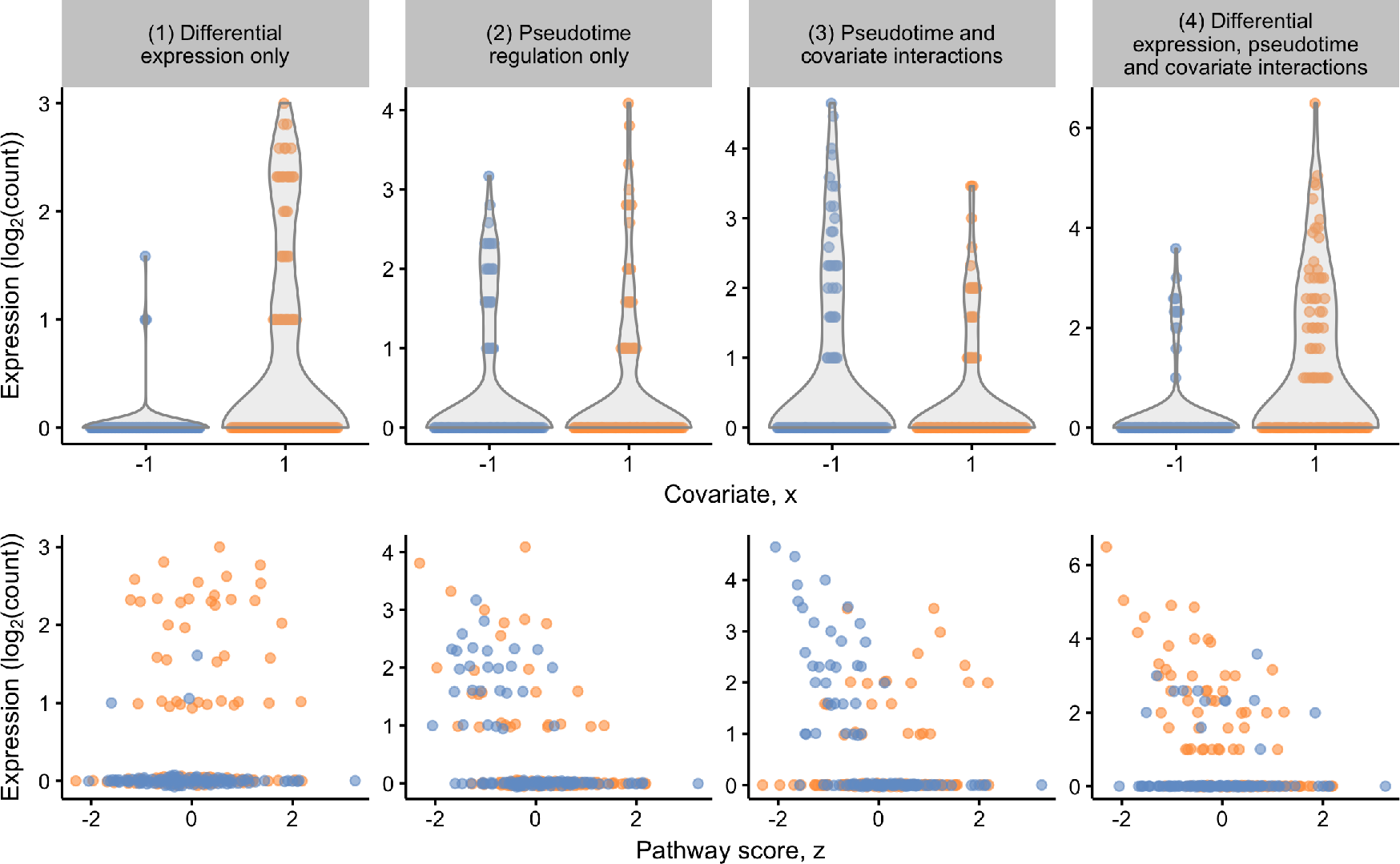
Four gene expression simulation scenarios were used: (1) differential expression only where the overall expression level for groups −1 and 1 differed but there is no dependence on pseudotime or pathway score, (2) pseudotime regulation only where the overall marginal distribution of expression values is identical between groups but expression changes with latent pathway score, (3) pseduotime and covariate interactions where the trajectory for each group differs over pathway score and (4) a complex scenario where differential expression and covariate-pseudotime interactions all exist.

**Supplementary Figure 2:**
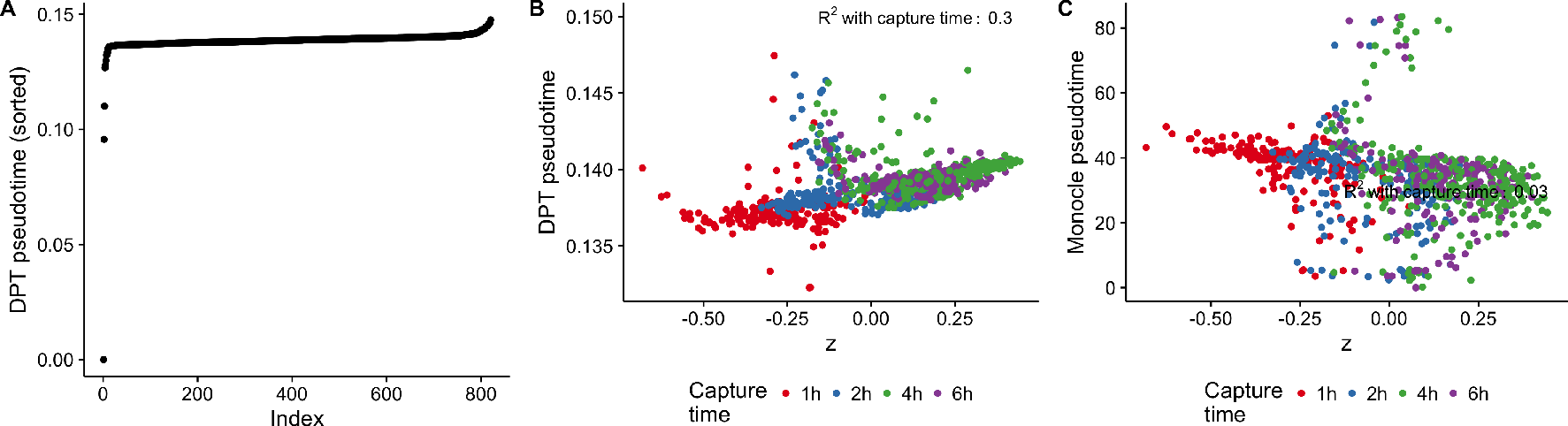
Performance of DPT and Monocle 2 on Shalek et al dataset. **A** Sorted DPT pseudotimes by index identifies three outlier cells. **B** Comparison of DPT pseudotimes to PhenoPath pathway score z. **C** Comparison of Monocle 2 pseudotimes to PhenoPath pathway score z.

**Supplementary Figure 3:**
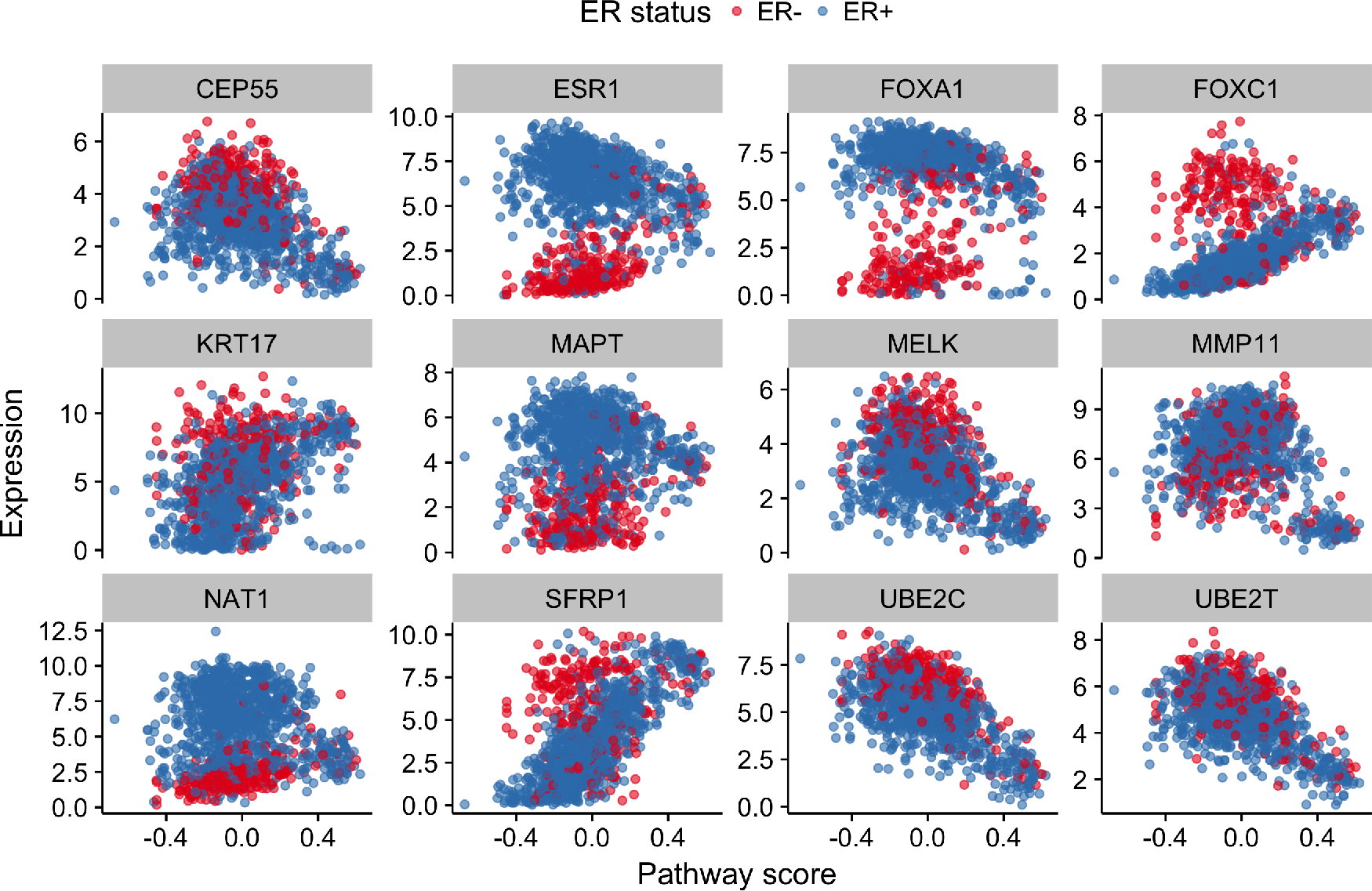
Pseudotemporally ordered gene expression trajectories for the TCGA Breast Cancer data for 12 breast cancer-associated genes.

**Supplementary Figure 4:**
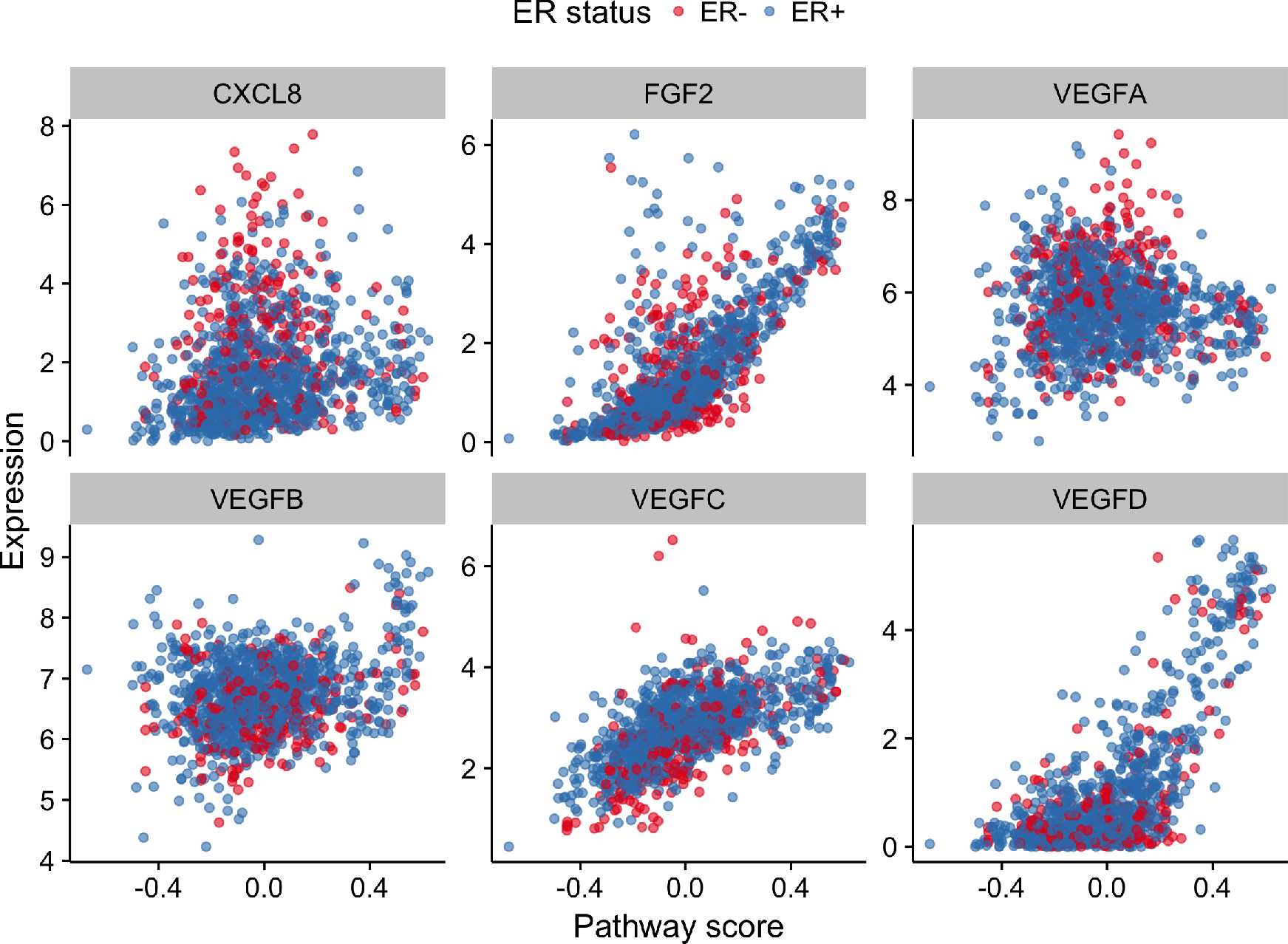
Pseudotemporally ordered gene expression trajectories for the TCGA Breast Cancer data for six angiogenesis-associated genes.

**Supplementary Figure 5:**
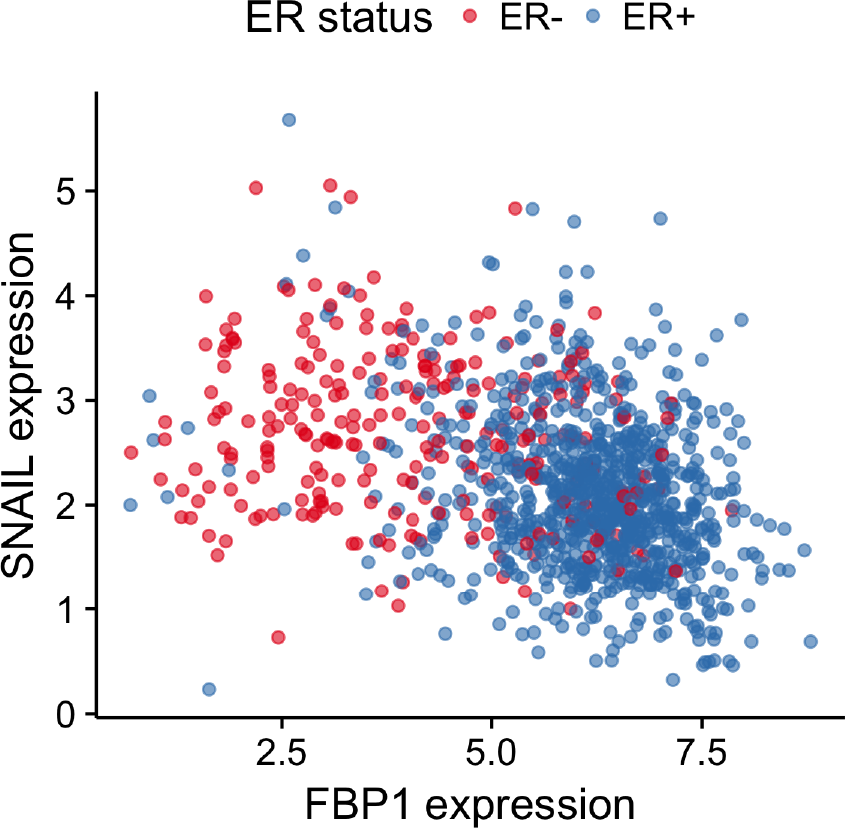
FBP1 expression is inversely correlated with Snail in ER-breast cancers but shows no dependence in ER+ breast cancers.

**Supplementary Figure 6:**
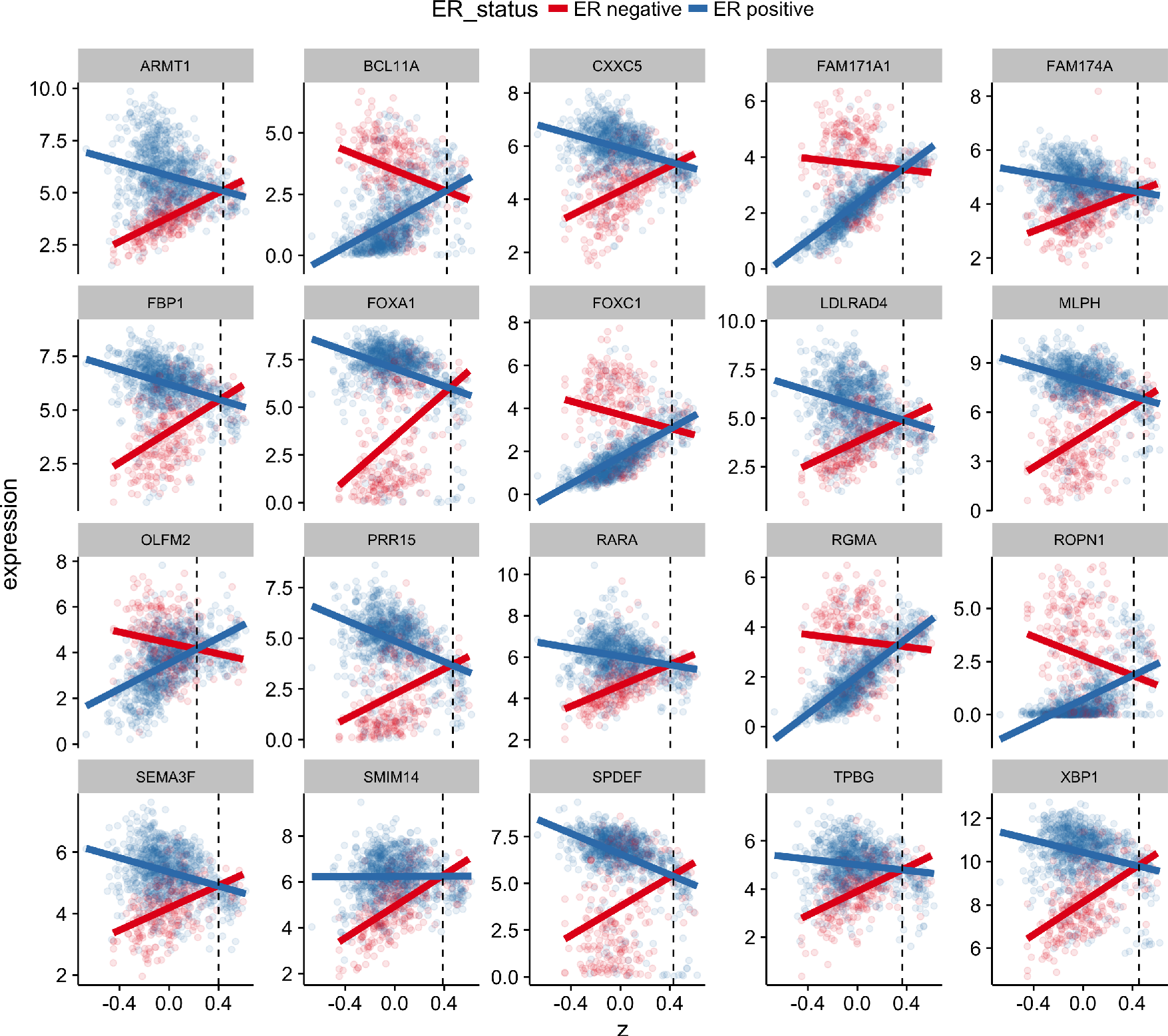
Expression of 20 genes with the largest interaction effects along the inferred pseudotemporal trajectory coloured by estrogen receptor status with linear fits as solid lines. The vertical dashed line indicates the crossover point.

